# Treatment of Murine Lupus with TIGIT-Ig

**DOI:** 10.1101/586321

**Authors:** Shuowu Liu, Lizhu Sun, Chuqi Wang, Yingshu Cui, Yuee Ling, Tian Li, Fangxing Lin, Wenyan Fu, Min Ding, Shuyi Zhang, Changhai Lei, Shi Hu

## Abstract

The TIGIT (T cell immunoreceptor with Ig and ITIM domains) protein is a co-inhibitory receptor that has been reported to suppress autoreactive T and B cells to trigger immunological tolerance. We generated a new recombinant protein by connecting the extracellular domain of murine TIGIT to the Fc region of the rat immunoglobulin IgG2a. The fusion protein was then characterized. The results suggested that among mice with lupus that were treated with the TIGIT-Ig fusion protein, the onset of proteinuria was delayed, serum concentrations of autoantibodies, such as antinuclear antibodies, were reduced without a decrease in the total IgG concentrations, and the survival rate was significantly increased compared to those of the controls. In conclusion, TIGIT-Ig administration showed promising results for both the prevention and treatment of autoimmune diseases in mice. This indicates that treatment with recombinant human TIGIT-Ig shows promise as an effective way to treat human autoimmune diseases.

## Introduction

Systemic lupus erythaematosus (SLE), a chronic inflammatory disease, is often difficult to diagnose because of its unpredictable patterns of recurrence and remission; SLE patients are predominantly female and especially of African, Asian or Hispanic origin[1]. Currently, SLE is incurable. Immunosuppressants are the mainstay of treatment, but their use increases the probability of infection in patients. SLE is characterized by the dysregulation of multiple arms of the immune system and by the production of hallmark autoantibodies. Genetic disorders, changes in female sex hormones, and environmental factors (such as sunlight, drugs, or smoking) are considered to be the causes of SLE. SLE exhibits many symptoms, including skin rashes, mouth ulcers, arthritis, neuropsychiatric syndrome, and blood cell damage. Serious complications such as kidney failure can further increase the morbidity and mortality of SLE. Autoantibodies bind to self-antigens and deposit the antigens into tissues, resulting in tissue damage, the loss of regulatory functions, and inflammatory cell infiltration, which leads to changes in the pathological structure of tissues and organs. Current treatments using steroids and cytotoxic drugs cannot cure SLE, but these treatments can inhibit the overactive immune system to prevent disease progression and organ damage. Therefore, there is a need for better therapeutic strategies that specifically target the pathogenic mechanisms while maintaining the protective function of the immune system. In addition, methods to predict disease onset and progression are important for better control of this complex disease [2].

In the last few decades, the understanding of SLE has increased greatly. As immune surveillance and immune activity are constantly monitored and balanced in the human body, current clinical immunotherapies purposefully regulate the immune system to keep it in order. T cell immunoreceptor with Ig and ITIM domains (TIGIT) is a novel immunomodulatory molecule and co-inhibitory receptor that was first identified by gene microarray screening for genes associated with immune responses [3]. TIGIT is expressed on effector and memory T cells, NK cells, and Treg cells and is known by different names, including Vstm3 [4] TIGIT [5], and WUCAM [6]. TIGIT is a type I transmembrane protein (N-terminal extracellular) composed of an immunoglobulin variable region (IgV)-like domain, a transmembrane region, and an ITIM domain. TIGIT can bind to two known ligands: CD155 (poliovirus receptor, PVR) and CD112 (PVRL2, nectin-2). CD155 is expressed on APCs, T cells, and various non-haematopoietic cell types, including cancer cells [7, 8]. There is two-way signalling between TIGIT and CD155; TIGIT stimulates CD155 on the DC surface, causing IL-10 secretion, and TIGIT is also stimulated by CD155 to inhibit the activation of NK cells and T cells. Similar to T cells, NK cells simultaneously express the co-stimulatory molecule CD226 and the co-suppressor molecule TIGIT, both of which bind to the same ligand, CD155; however, the TIGIT co-suppression signal is usually dominant under normal conditions [9].

Previous studies have shown that the CD28-B7.1/B7.2 co-stimulatory signal of T cells can be reduced and blocked by CTLA4-Ig[10], which is a fusion protein consisting of the extracellular domain of CTLA-4 and the Fc segment of IgG1. Abatacept is a first-generation CTLA4-Ig co-stimulation blocker for the treatment of rheumatoid arthritis[11]. In an SLE mouse model that was treated with CTLA4-Ig, the onset of SLE was delayed, autoantibody production was reduced, proteinuria was reduced, and survival was prolonged compared to those of the controls[12]. The CTLA-4/CD28-CD80-CD86 and TIGIT/CD226-CD155-CD112 signalling pathways are similar; the homologous molecules CD155 and CD112 can stimulate the co-suppressor molecule TIGIT, inhibiting T cell and NK cell activation, while the corresponding co-stimulatory molecule CD226 promotes activation; additionally, the homologous molecules CD80 and CD86 can bind to the co-suppressor molecule CTLA-4, inhibiting T cell activation, while binding to the corresponding co-stimulatory molecule CD28, which promotes the activation of T cells. The difference between these two pathways is that CD80/CD86 is mainly expressed in APCs, while the expression of CD155 is very extensive, and, while CD155 is not expressed by DCs, CD155 is expressed by a variety of non-professional APCs (such as the vascular endothelium, fibroblasts, and tumour cells) [13, 14]. When autoimmune diseases occur, the tissue that is infiltrated by T cells is contains mainly non-professional APCs, suggesting that the mechanism of the TIGIT-CD226-CD155-CD112 network in autoimmune diseases, tumour immunity and other sources of inflammation needs to be further studied[15].

Here, we introduced a new approach to treat lupus-prone (NZB/NZW) F1 mice by using TIGIT-Ig. Delayed proteinuria, a milder inflammatory response, an improved survival rate and a reduction in the production of autoantibodies against dsDNA and histones compared to those of the controls were observed, suggesting that TIGIT-Ig may be a promising clinical treatment.

## Material and methods

### Generation of fusion proteins

Based on previous studies [16–18], the sequences of the extracellular domains (ECDs) of TIGIT and CTLA4 were ligated to the Fc segment of murine IG2a, which is the gene that codes for the hinge-CH2-CH3 segment, to construct a recombinant plasmid. The FreeStyle 293 expression system (Invitrogen) was used in our study, and the recombinant protein was obtained according to the methods used in a previous study [19] and then immediately purified using protein A-Sepharose and the harvested cell culture supernatant. The concentration and purity of the fusion protein were determined by measuring the UV absorbance at a wavelength of 280 nm and by polyacrylamide gel electrophoresis, respectively.

### Affinity Measurement

We immobilized an anti-murine Fc polyclonal antibody (Jackson ImmunoResearch Europe Ltd.) on a CM5 chip (~150 RU) using standard amine-coupling chemistry and then injected CD155ECD or B7 (12.5 nM~200 nM) using a previously reported method to capture the fusion proteins. The binding response was corrected by subtracting the RU from the blank flow cells. We used the surface plasmon resonance (SPR) method with a BIAcore-2000 to measure the monovalent binding affinity of the fusion protein and performed kinetic analysis using a 1:1 Langmuir model that simultaneously fit k_a_ and k_b_.

### Pharmacokinetics

We used female C.B-17SCID mice to determine the pharmacokinetic profile of the fusion protein. Eight-week-old mice were administered the fusion protein at a dose of 1 mg/kg body weight by tail vein injection. Mice were divided into 15 groups, corresponding to day 1 to day 15. Blood was collected from the septum in heparin-containing tubes and then centrifuged to remove blood cells and to obtain plasma samples. The serum concentration of the fusion protein was determined by competitive ELISA.

### Mice and treatment

We used female NZB/NZW F1 mice to conduct our experiments. The mice were purchased from Harlan Winkelmann (Borchen, Germany) when they were 5-7 weeks old. Sixteen-week-old mice (prophylactic, n=30) or 33-week-old mice (therapeutic, n=40) were separately intraperitoneally injected for 10 weeks. Mice in each group were treated with 100 μg of two kinds of fusion proteins or with control IgG twice a week. All experiments involving animals were conducted in accordance with the guidelines and regulations for animal use and care of the Experimental Animal Center of the Chinese Academy of Sciences (Beijing, China).

### Quantification of IgG and autoantibodies

We immobilized goat anti-mouse IgG (Southern Biotech) to Maxisorp plates (Nunc, Roskilde, Denmark) and blocked the plates with 2% FCS using a method reported by Charlotte Starke[20]. Bound mouse IgG was detected with HRP-conjugated goat anti-mouse IgG (Southern Biotech), and the total IgG concentration was calculated with mouse IgG standards (Jackson, West Grove, USA). Then, we pre-incubated the plates with poly-L-lysine (Sigma, Saint Louis, MO) or high-purity histone (Roche Diagnostics, Mannheim, Germany), dissolved them overnight at 4 °C, and then washed and incubated the plates with calf thymus DNA (Sigma) overnight at 4 °C, which was used to measure anti-dsDNA or anti-histone IgG. Antibody titres were calculated by performing high (S1) and low (S2) ELISAs against the corresponding antibody antigens based on the two reference sera, which were used as internal standards in each assay. The antibody titres were taken as relative units, with the cOD value of S1 subtracted from the cOD value of S2 and the result divided by 100 [1 unit = (cODS1 - c0DS2)/100].

### Quantification of antibody-secreting cells

Similarly, we pre-coated 96-well multi-screen plates with 1 μg/ml goat anti-mouse IgG (Southern Biotech) or 20 μg/ml poly-L-lysine (Sigma) together with 20 μg/ml calf thymus DNA (Sigma) (Millipore, Billerica, USA) and then used 2% FCS in PBS to block the plates[20]. Single-cell suspensions of lymphocyte M (Cedarlane Laboratories Ltd., Hornby, Canada)-purified PBMCs at 0.25×10^5^ cells/well, bone marrow cells (2 femurs) at 1×10^5^ cells/well and splenic cells at 0.5×10^5^ cells/well were added to the plates and incubated for 1 hour at 37 °C. The plates were washed, and HRP-conjugated goat anti-mouse IgG (Jackson ImmunoResearch Laboratories) was added to the plates and incubated at 20 °C for 1 hour.

### Proteinuria

Urine was collected weekly and subjected to standard semi-quantitative tests to determine the urinary protein levels. We measured the urine protein levels using Bayer Multistix test strips (Bayer, Fernwald, Germany). As outlined in the instructions, slightly referred to 15-20 mg/d1 albumin, one plus referred to 30 mg/dl albumin, two plus referred to 100 mg/dl albumin, three plus referred to 300 mg/dl albumin, five plus referred to 2,000 mg/dl albumin, and nephritis referred to proteinuria of ≥300 mg/dl albumin.

### Flow cytometry

Whole blood cells stained with anti-CD4-FITC and anti-CD8-PE antibodies (BD Biosciences, Heidelberg, Germany) were detected with a FACSCalibur flow cytometer and then analysed with Cell Quest software (BD Biosciences).

### Histopathology

Mouse kidneys were obtained, fixed with paraformaldehyde, embedded in paraffin, and cut into 4-μm sections. The sections were stained with haematoxylin and eosin (H&E) and then observed and analysed with light microscopy (Nikon, Tokyo, Japan). Next, the total glomerulosclerosis damage and tubulointerstitial damage were assessed as previously reported using a damage score that was calculated with the equation [(2×N glomeruli with +2 score) + (3×N glomeruli with +3 score) + (4×N glomeruli with +4 score) + (1×N glomeruli with +1 score)]/total number of glomeruli examined to assess total glomerulosclerosis damage; the extent of the damaged tubulointerstitial area in the renal cortex was classified with 0 = normal; grade +1 = <10%; grade +2 = 10–25%; grade +3 = 25–50%; grade +4 = 50–75%; and grade +5 = 75–100%)[21].

### Statistical analysis

Proteinuria and survival are displayed in the Kaplan-Meier plots. Statistically significant differences were determined by a Student’s unpaired t test unless otherwise indicated. P < 0.05 was considered significant.

## Results

### TIGIT-Ig fusion protein

To investigate the therapeutic potential of TIGIT, we constructed and generated a fusion protein (TIGIT-Ig) consisting of the extracellular domain of murine TIGIT linked to the murine IgG2a chain. The expression and purification methods were described in previous reports [21]. As previously reported [22], the affinity of the fusion protein for CD155 was determined with BIAcore binding assays (Fig S1).

Compared with the well-studied Fc fusion protein CTLA-4-Ig, the Fc fusion protein TIGIT-Ig had a similar denaturation temperature and thus exhibited IgG-like stability. The lowest concentration (< 2%) of high molecular weight and low molecular weight products was observed after 3 weeks of storage at 40 °C at a 1 mg/mL concentration. Mice were treated separately with a single intravenous dose of TIGIT-Ig and CTLA4-Ig to measure their pharmacokinetic (PK) parameters, and the serum concentrations of TIGIT-Ig and CTLA4-Ig were determined by ELISA. The results showed that the main PK parameters of TIGIT-Ig and CTLA4-Ig were very similar in mice and demonstrated the high stability of the TIGIT-Ig fusion protein. The experimental data are summarized in Table S1.

### TIGIT-Ig notably inhibits the production of autoantibodies in lupus-prone F1 mice

To test the ability of TIGIT-Ig to provide protection and the efficacy of TIGIT-Ig, we first intraperitoneally (IP)administered 100 μg TIGIT-Ig to female lupus-prone (NZB/NZW) F1 mice (n = 30) twice a week for 10 weeks. Then, we determined the serum concentration of TIGIT-Ig by ELISA, and we established a dosing regimen that maintained a serum concentration of no less than 5 μg/ml. During the treatment period, we monitored the concentrations of the total IgG and anti-histone and anti-dsDNA antibodies because SLE is most likely correlated with anti-dsDNA antibody production or anti-histone autoantibody production.

IgG autoantibodies against dsDNA and histones were detectable at 16 weeks in all treatment groups (Fig. 1A and B). In the control group, the serum titres of the anti-dsDNA autoantibodies increased rapidly when the mice were 25 weeks old (Fig. 1A). Interestingly, the 40-week-old mice showed only a small increase in the anti-dsDNA autoantibody levels after treatment with the TIGIT-Ig fusion protein, in contrast to the control mice whose anti-dsDNA autoantibody levels greatly increased. Additionally, we did not detect increased levels of the anti-dsDNA autoantibodies in the TIGIT-Ig treatment group until 15 weeks later, 29 weeks after the last application of TIGIT-Ig. Although TIGIT-Ig treatment did not seem to delay the development of anti-histone IgG, the autoantibody production was significantly inhibited at week 40 compared to that of the control groups.

**Figure 1.**
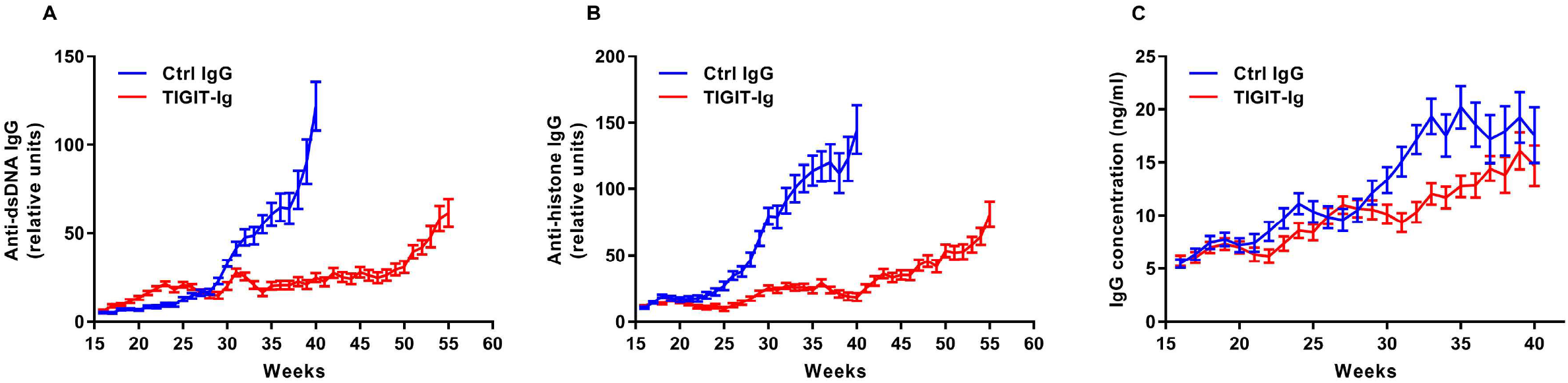
Autoantibody production was reduced in female NZB/NZW F1 mice with lupus that were treated with TIGIT-Ig. Serum concentrations of (A) anti-dsDNA IgG, (B) anti-histone IgG and (C) total IgG antibodies were determined by ELISA in two groups of F1 mice with lupus that were treated with different proteins. Relative units were calculated using the internal reference sera in each assay. Data represent mean values+SDs.

### Treatment with TIGIT-Ig markedly delayed the development of proteinuria

Mice that were prophylactically treated with TIGIT-Ig prior to the clinical onset of lupus, which started at week 16, showed a significant delay in proteinuria compared to that of the control mice treated with the control IgG (P < 0.05) (Fig. 2A). At the age of 36 weeks, all control mice developed proteinuria, whereas only 23.33% of the mice treated with TIGIT-Ig (P < 0.001) developed proteinuria. At 52 weeks, all TIGIT-Ig-treated mice had proteinuria. The TIGIT-Ig significantly lowered blood urea nitrogen (BUN), creatinine, and blood pressure compared to those of the controls, as shown in Table S2.

**Figure 2.**
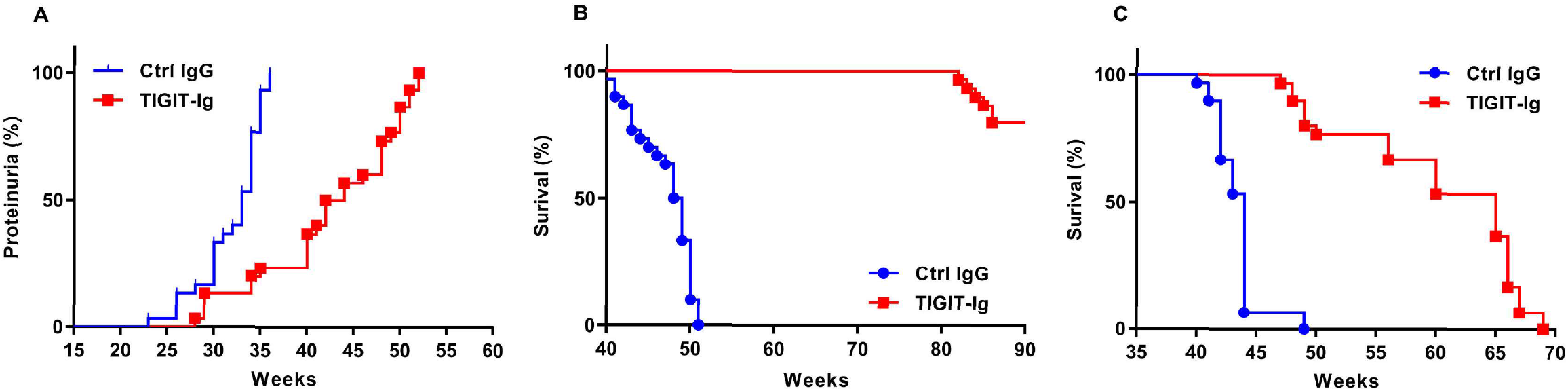
The incidence of proteinuria was low and the survival rate was high in NZB/NZW F1 mice with lupus. **(A)** Comparison of the incidence of proteinuria in mice treated with TIGIT-Ig and mice treated with IgG. **(B)** Comparison of the survival of mice treated with the fusion protein and mice treated with IgG. (C) Comparison of the survival of mice with active autoimmune disease treated with the fusion protein and mice treated with IgG.

### Effect of TIGIT-Ig treatment on the survival rate and the progression of disease in pre-disease female NZB/NZW F1 mice

Figure 2A and 2B show that compared with the control group, the TIGIT-Ig group exhibited a dramatically extended survival time and benefitted from a reduction in the levels of urine protein and/or autoantibody production (P < 0.001). Excitingly, all TIGIT-Ig treated mice survived to the age of 50 weeks, while the control mice had all died by that point.

The effects of TIGIT-Ig treatment were maintained, although the administration of TIGIT-Ig stopped when the mice were 26-weeks-old. At week 84, after the cessation of treatment, only 23.33% of the mice treated with TIGIT-Ig had died, while all of the control mice died. After treatment, the serum concentration of TIGIT-Ig decreased to 0.1 μg/ml or less within 4 weeks. Therefore, despite the decrease in its serum concentration, the beneficial effects of TIGIT-Ig remained (Fig. 2B).

### The potential of TIGIT-Ig treatment to influence peripheral blood lymphocyte subsets

Given the significant impact of TIGIT on T cell inhibition and tolerance, we analysed the peripheral blood lymphocytes of NZB/NZW F1 mice immediately after the treatment period of TIGIT-Ig administration to study the potential effects of TIGIT-Ig on circulating T lymphocytes in mice. We found that at the end of the treatment period, there was no marked difference in the percentage of peripheral CD4^+^ and CD8^+^ T cells (Fig. 3A) between the TIGIT-Ig-treated mice and the control mice, as analysed by flow cytometry. Murine IgG2a can bind to its complement, so TIGIT-Ig may effectively reduce the number of activated T cells through the complement pathway. However, our study showed that the absolute number of circulating T cells was not significantly altered by TIGIT-Ig treatment (Fig. 3A) compared to that of the controls. Our data also showed that compared to those of the controls, there was a minor increase in CD45RO expression in both CD4^+^ and CD8^+^ T cells with TIGIT-Ig treatment (Fig. 3B), while no alterations in the expression of the analysed surface marker CD 69 were observed (Fig. 3E). Moreover, compared to that of the controls, the CD161 levels were increased on both of the CD4^+^ T cell subsets in CSF, particularly on CD8^+^ T cells (Fig. 3D). Compared to those of the controls, the FoxP3^+^ cell population was not altered by TIGIT-Ig treatment (Fig. 4A), but the CD40L expression levels in CD4^+^ T cells was reduced more with TIGIT-Ig treatment than without the treatment (Fig. 4B).

**Figure 3.**
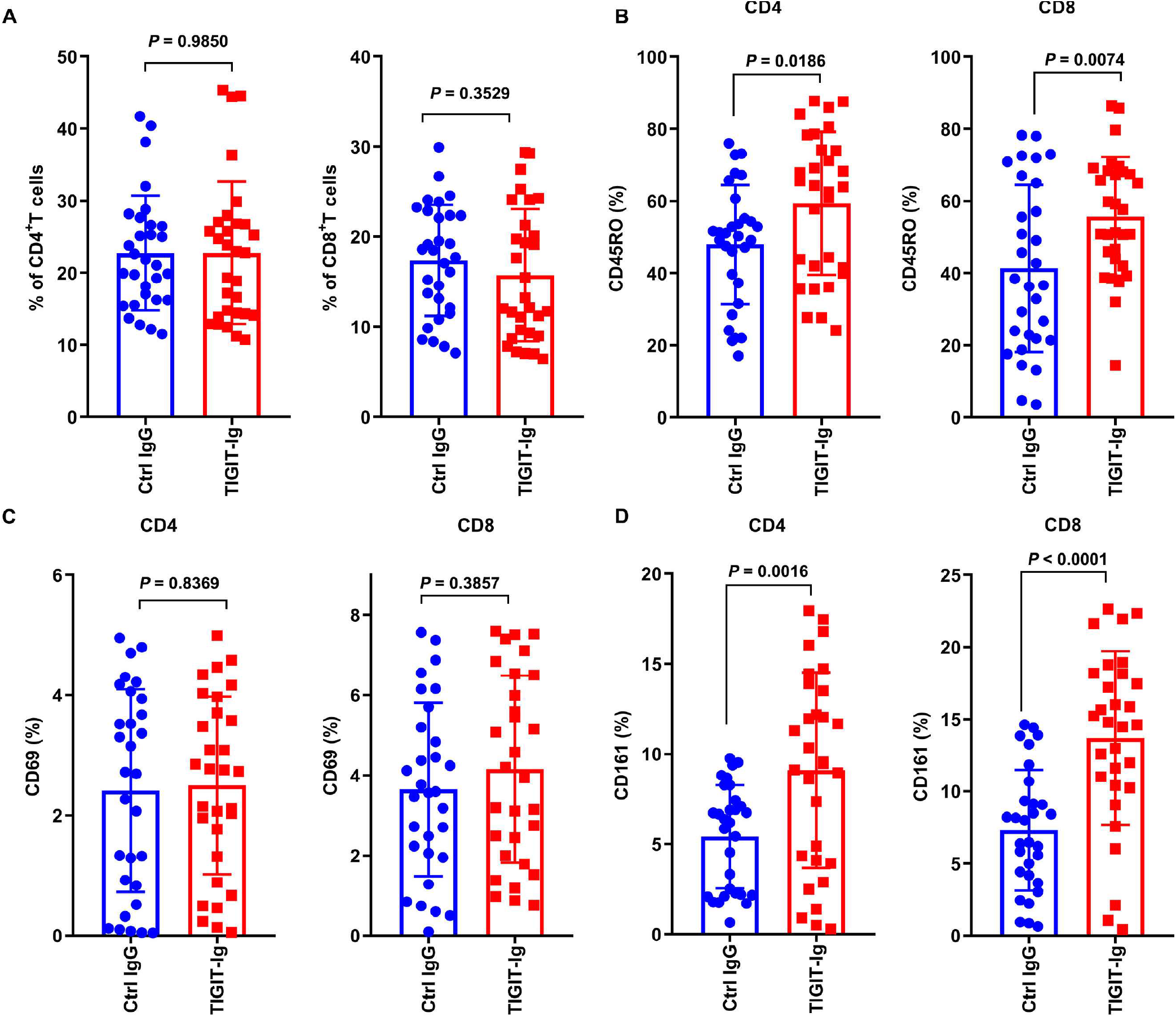
The percentage of peripheral T lymphocytes was not notably altered by the fusion protein among TIGIT-Ig-treated mice. For FACS analysis, whole blood was gated for lymphocytes with both the forward and side scatter. **(A)** Percentages of CD4^+^ cells **and** CD8^+^ cells. Both cell types were analysed by flow cytometry of the peripheral blood of the fusion protein- or IgG-treated mice. Each data point on the charts represents an individual mouse. Horizontal bars indicate the mean value. Cells within the CD4^+^ and CD8^+^ T cell subsets expressing CD45RO (B), CD69 (C), and CD161(D) were measured.

**Figure 4.**
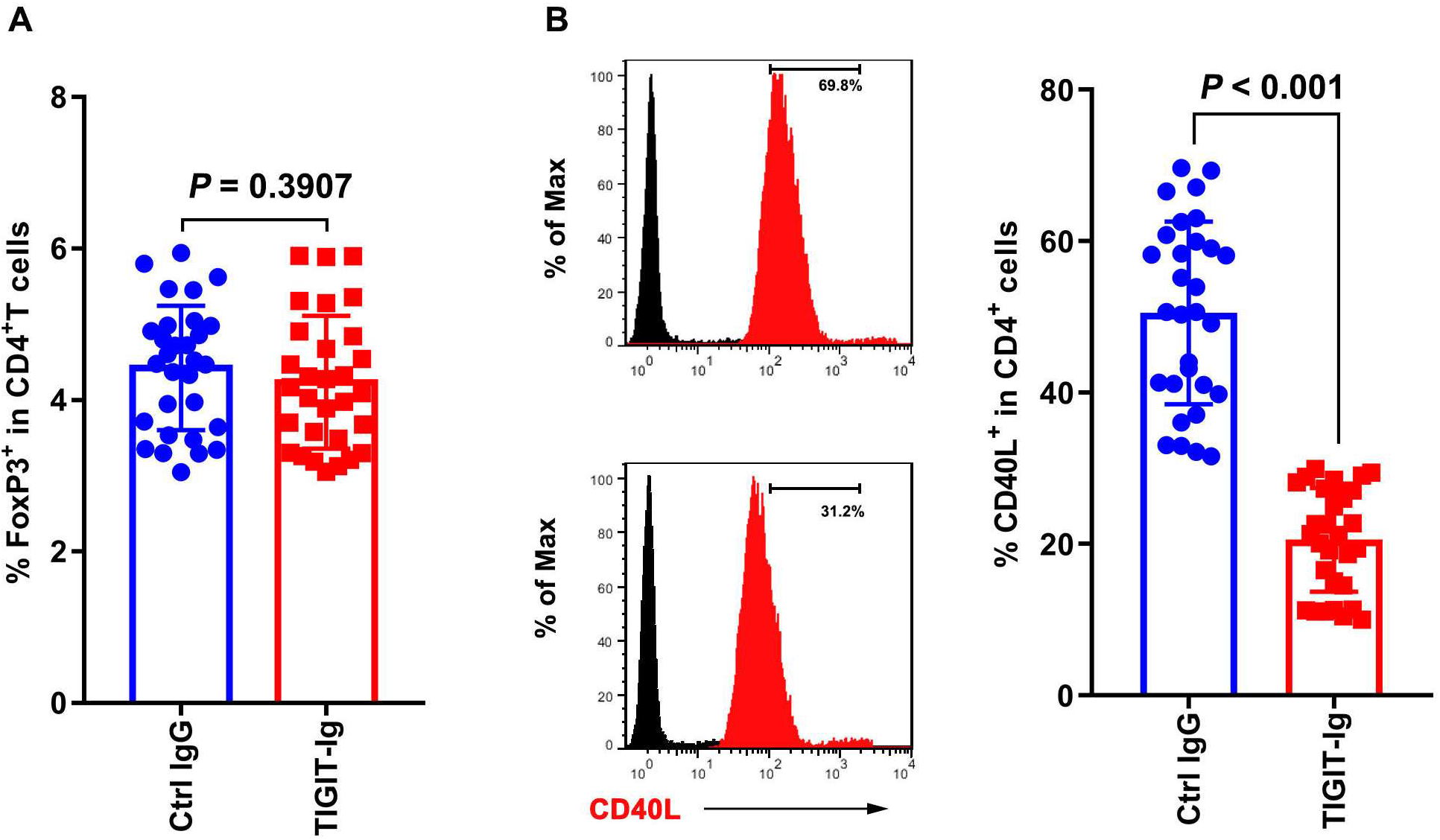
The circulating cytokine milieu was not significantly changed in the fusion protein-treated NZB/NZW F1 mice. (A) Percentages of CD4^+^/FoxP3^+^ positive cells. (B) Percentages of CD40L^+^ cells. Both cell types were measured with flow cytometry of the peripheral blood of the fusion protein- or IgG-treated mice. Each data point represents an individual mouse.

### Kidney pathology analysis after TIGIT-Ig administration in lupus-prone mice

Histopathological kidney sections were stained with H&E and then observed and analysed with light microscopy after the prophylactic treatment study had ended. The results were in agreement with those shown by the previously presented data, and mice treated with TIGIT-Ig showed only minor glomerular lesions (Fig. 5A, B, C and D) compared to that in the control group. In contrast, significant glomerular changes, such as glomerular hypercellularity, renal interstitial fibrosis with inflammatory cell infiltration, mesangial expansion, proliferation of the mesangial matrix, and global sclerosis, were observed in mice treated with IgG (Fig. 5E and F). We also clearly observed many swollen and necrotic tubule epithelial cells, cell debris and protein deposits in the TIGIT-IG group. However, we noticed a marked decrease in tubular proteinosis in the TIGIT-Ig-treated mice.

**Figure 5.**
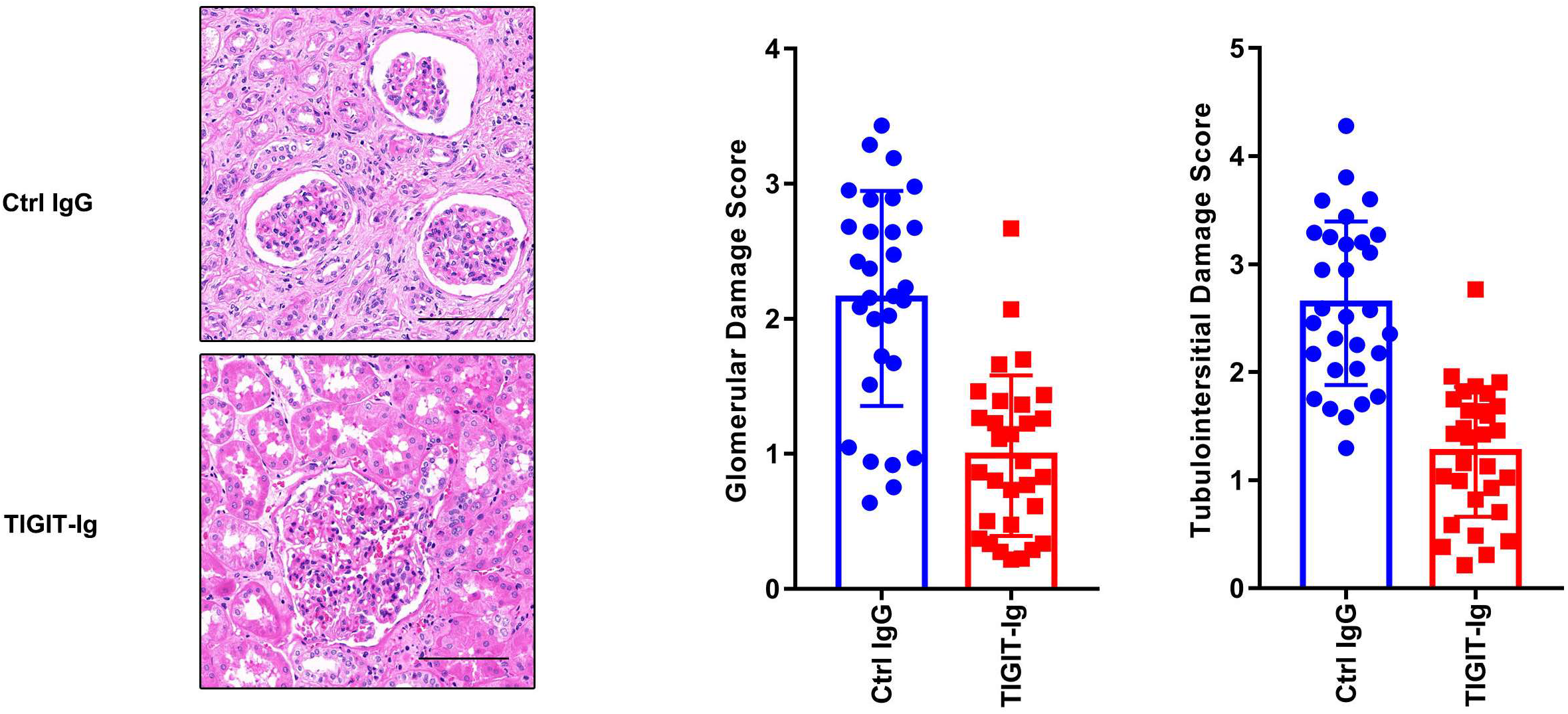
The severity of kidney pathology was reduced by TIGIT-Ig treatment. (A) Control mice developed glomerular hypercellularity, tubulointerstitial inflammation, and some epithelial cell necrosis and proliferation in the renal proximal convoluted tubules. (B) A marked decrease in glomerular hypercellularity, tubular proteinosis, and interstitial congestion were observed in mice treated with TIGIT-Ig compared to those in the controls. (C) Mice treated with TIGIT-Ig showed mild glomerular damage compared to that in the control group. Glomerular damage scores for different treatments are presented. (D) A significant reduction in tubulointerstitial damage was also observed in the TIGIT-Ig group compared to that in the control group. Tubulointerstitial damage scores for the different treatments are presented.

### Reduced number of splenic and peripheral blood cells that secrete IgG in treated mice with lupus

We used two additional groups of NZB/W F1 mice to administer the fusion proteins for 10 weeks starting at 16 weeks of age to perform in vivo testing (n = 10) to determine the total amount of IgG and antibodies at 36 weeks of age and to analyse whether TIGIT-Ig treatment effectively inhibited (autologous) antibody-secreting cells (ASCs).

Then, we used ELISPOT to analyse the number of IgG-secreting cells present in the bone marrow, spleen and among the peripheral blood mononuclear cells (PBMCs). The number of IgG-secreting cells in the TIGIT-Ig-treated mice was reduced in comparison with that in the control group (Fig. 6A, B and C). In addition, we noticed that the number of anti-dsDNA IgG-secreting cells, which are autoreactive cells present in the peripheral blood, bone marrow and spleen, of mice treated with TIGIT-Ig was lower than in control mice (Fig. 6D, E and F). Therefore, from the abovementioned studies, we concluded that the proliferation, survival and/or migration of splenic and peripheral ASCs may be affected by TIGIT-Ig treatment. When bone marrow-derived dendritic cells (BMDCs) were differentiated from both the control and TIGIT-Ig-treated groups and analysed by flow cytometry, the percentage of CD11c^+^ and CD11c^+^CD11b^+^ cells obtained from the TIGIT-Ig-treated mice was significantly lower than that in control mice (Fig. 7A). Additionally, the percentages of GC B cell plasma cells and Tfh cells (Fig. 7B) were significantly reduced in mice treated with TIGIT-Ig compared with those in control mice.

**Figure 6.**
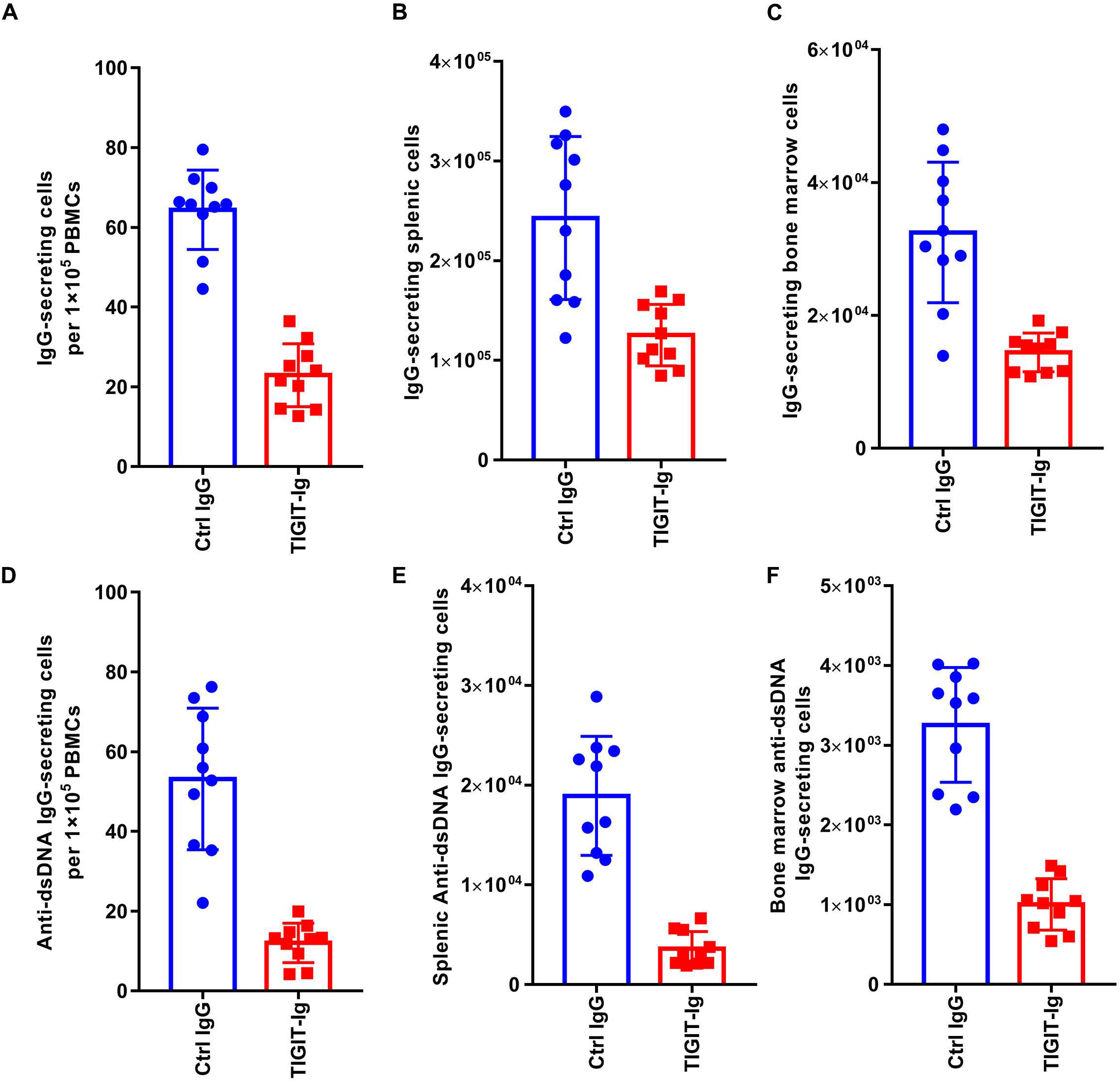
Total IgG antibody-secreting cells (ASCs) were reduced in the spleen, bone marrow and peripheral blood of the fusion protein-treated mice with lupus. ELISPOT results showed the number of IgG ASCs detected in the isolated PBMCs **(A)**, spleen **(B)** and bone marrow **(C)** of the IgG- or fusion protein-treated 33-week-old NZB/W F1 mice (n = 10 in each group). Autoreactive anti-dsDNA IgG-secreting cells in PBMCs **(D)**, spleen **(E)** and bone marrow **(F)** are shown. Horizontal bars indicate the mean values+SDs. *p < 0.01.

**Figure 7.**
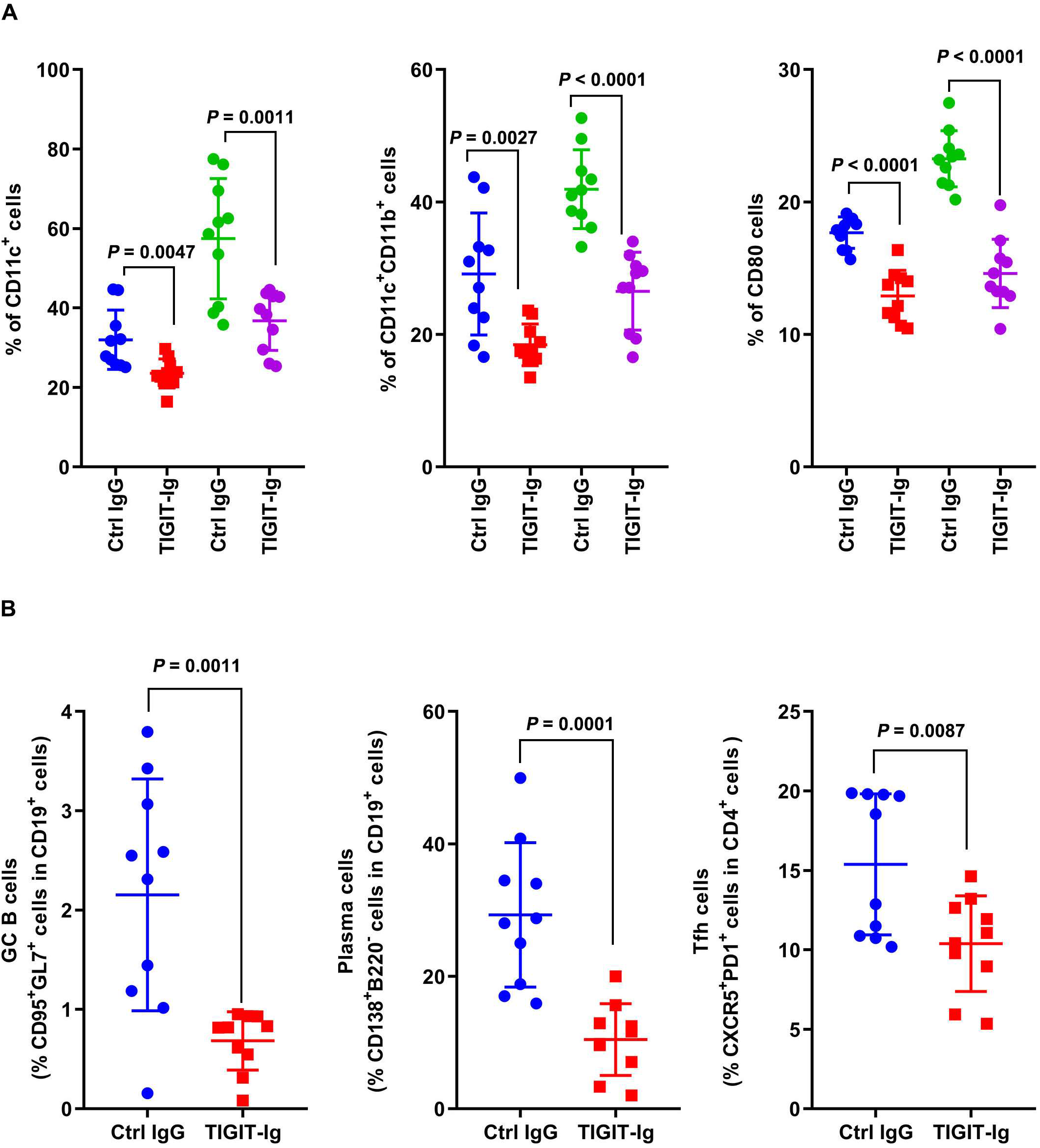
BMDCs and GC lymphocytes were affected by TIGIT-IG treatment. BMDCs from mice that were stimulated with LPS for 24 hrs. **(A)** Percentages of CD11c^+^-, **(B)** CD11b^+^CD11c^+^-, and **(C)** CD80^+^-expressing BMDCs. **(D–F)** The frequencies of GC B cells, CD138^+^ plasma cells, and Tfh cells were analysed by flow cytometry. The percentages of GC B cells (gated on the CD19^+^ population), CD138^+^ cells, and Tfh cells (gated on the CD4^+^ population) are shown.

### Therapeutic effects of TIGIT-Ig administration in lupus-susceptible mice

To investigate the therapeutic effects of TIGIT-Ig on active autoimmune disease, we monitored female (NZB/NZW) F1 mice up to an age of 35 weeks and selected those with established progressive lupus whose urinary protein levels were not lower than 300 mg/dL for in vivo experiments. These mice were divided into two groups with 30 mice in each group and treated with 100 μg TIGIT-Ig or with control IgG intraperitoneally (IP) three times a week for 10 weeks in the late stage of the disease.

After 10 weeks of therapy, only 0.06% of the control mice had survived, whereas all of the mice in the therapy group (P < 0.01) had survived. Fourteen weeks after the termination of treatment, all mice in the control group had died, while 80% of the mice in the therapy group (P < 0.0001) survived to an age of 49 weeks (Fig. 2C). Therefore, TIGIT-Ig has an inhibitory effect on established pathological immune responses.

## Discussion

In this study, we treated NZB/NZW F1 mice for several months with TIGIT-Ig, a fusion protein containing the murine TIGIT-ECD linked to the murine IgG2a chain and demonstrated for the first time that it could be used as a potential therapeutic agent. The results of this study showed that after treatment, the mice had decreased proteinuria, reduced autoantibody production, and prolonged survival. In addition, the number of IgG-secreting cells and anti-dsDNA-secreting cells in the spleen, bone marrow and peripheral blood were also significantly reduced in the TIGIT-Ig-treated group compared to those in the control group. Histological studies also showed that TIGIT-Ig significantly inhibited renal tubular proteinosis in mice. Together, these studies demonstrate the effectiveness of TIGIT therapy. More importantly, the TIGIT-Ig fusion protein therapy did not cause lymphocyte depletion, and no neutralizing anti-TIGIT-Ig antibodies were detected in the treated mice, even at the end of the treatment period. We know that biological agents such as cytokine-targeting inhibitors and targeted immune cell antagonists can cause a host immune response that includes reduced efficacy or even toxicity during the treatment of chronic autoimmune diseases. Our study indicated that the TIGIT-Ig fusion therapy overcame some of these problems.

In recent years, a growing number of co-suppressor receptors that inhibit T cells and other immune responses have been discovered. In 2006, Ahmed’s team demonstrated that the co-inhibitory receptor PD-1 can limit CD8^+^ T cell responses, and therefore, it may be used as an important checkpoint inhibitor [23]. It was later discovered that T cell co-suppressive factors such as CTLA-4, LAG-3 and TIM-3, together with PD-1, can gradually deplete the response of protective CD8^+^ T cells [24] in the case of chronic infectious diseases and other persistent immune-stimulated non-infectious diseases such as cancer. TIGIT has recently been found to be highly associated with T cell tumour infiltration [25], along with PD-1 and CTLA-4. Studies have shown that TIGIT can regulate immune responses in a variety of ways, including intracellular and extrinsic pathways and that TIGIT is as highly associated with T cell tumour infiltration, similar to PD-1 and CTLA-4 [25]. There have been many recent advances in the understanding of the mechanism of TIGIT-induced suppression of the immune response. First, a study found that CD8^+^ TILs in TIGIT^-/-^ mice exhibited high cytotoxicity and increased proliferation [26]. CD8^+^ T cells in TIGIT knockdown patients with acute myeloid leukaemia also showed the same results [27]. In addition, studies have shown that the synergistic blocking of TIGIT and PD-1 in mice with colon cancer can induce the increased secretion of cytokines, such as interferon-γ and tumour necrosis factor-α [24], the antigen-specific proliferation of CD8^+^ T cells among TILs; additionally, CD8^+^ T cells among the peripheral blood mononuclear cells and TILs can be degranulated in melanoma patients [28]. Thus, these findings all demonstrate the role of TIGIT in inhibiting the proliferation of CD8^+^ T cells and in exerting effects on CD8^+^ T cells.

Moreover, recent reports in cancer research have also provided insight into the mechanisms underlying TIGIT. A study found that TIGIT in Treg cells is also responsible for the inhibition of immune responses. By suppressing the antitumour immune response, TIGIT can selectively inhibit the antitumour type 1 Th17 response in Treg cells, but Treg cells retain the type 2 response (IL-4), promoting the differentiation of type 2 tumour-associated macrophages (TAM2); these cells inhibit T cell responses by secreting inhibitory cytokines such as IL-10 and TGF-β as well as enzymes such as arginase-1 and indoleamine 2,3-dioxygenase (IDO) to deplete the nutrients in the tumour microenvironment [29]. Studies have shown that IL-10, which is an important immunosuppressive cytokine that promotes DC tolerance and plays an important role in the tumour microenvironment, can be produced in large quantities by TIGIT^+^ Treg cells in the tumour environment [26]. Moreover, TIGIT+ Treg cells can also regulate the DC phenotype through the TIGIT-CD155 signalling pathway, inhibiting the activation and effector functions of CD8+ T cells and NK cells. However, the in vivo mechanisms of TIGIT-mediated NK cell-induced anti-metastasis and antitumour immune surveillance need to be studied further [30].

In conclusion, TIGIT-Ig-treated mice exhibited reduced autoantibody production, which alleviated the disease response and prolonged survival compared to those of the controls. TIGIT-Ig is expected to be an effective treatment for autoimmune diseases.

## Acknowledgments

We thank Y. Bao and Z. Zou (Changzheng Hospital Translational Medicine Center) for generous help of the research. This study was financially supported by the National Natural Science Foundation of China (grant no. 81773261 and 81602690); Shanghai Sailing Program (19YF1438600); Shanghai Rising-Star Program (19QA1411400), Shanghai Chenguang Program (17CG35); Military Medicine Special Grant of Second Military Medical University (grant no.2017JS01); a General Financial Grant from the China Postdoctoral Science Foundation (grant no. 2016M593006), and postdoctoral scientific research funds of Second Military Medical University.

