## Supplementary for "Treatment of Murine Lupus with TIGIT-Ig"

**Contents**	Page

Supplementary Figures	2

Figure S1. Binding of Fusion proteins to CD155 and B7.	2

Supplementary Tables	3

Table S1. Selected analytical data and pharmacokinetic parameters of recombinant fusion proteins in mice.	3

Table S2. Biochemical characteristics after 10 weeks of treatment	3

### Supplementary Figures

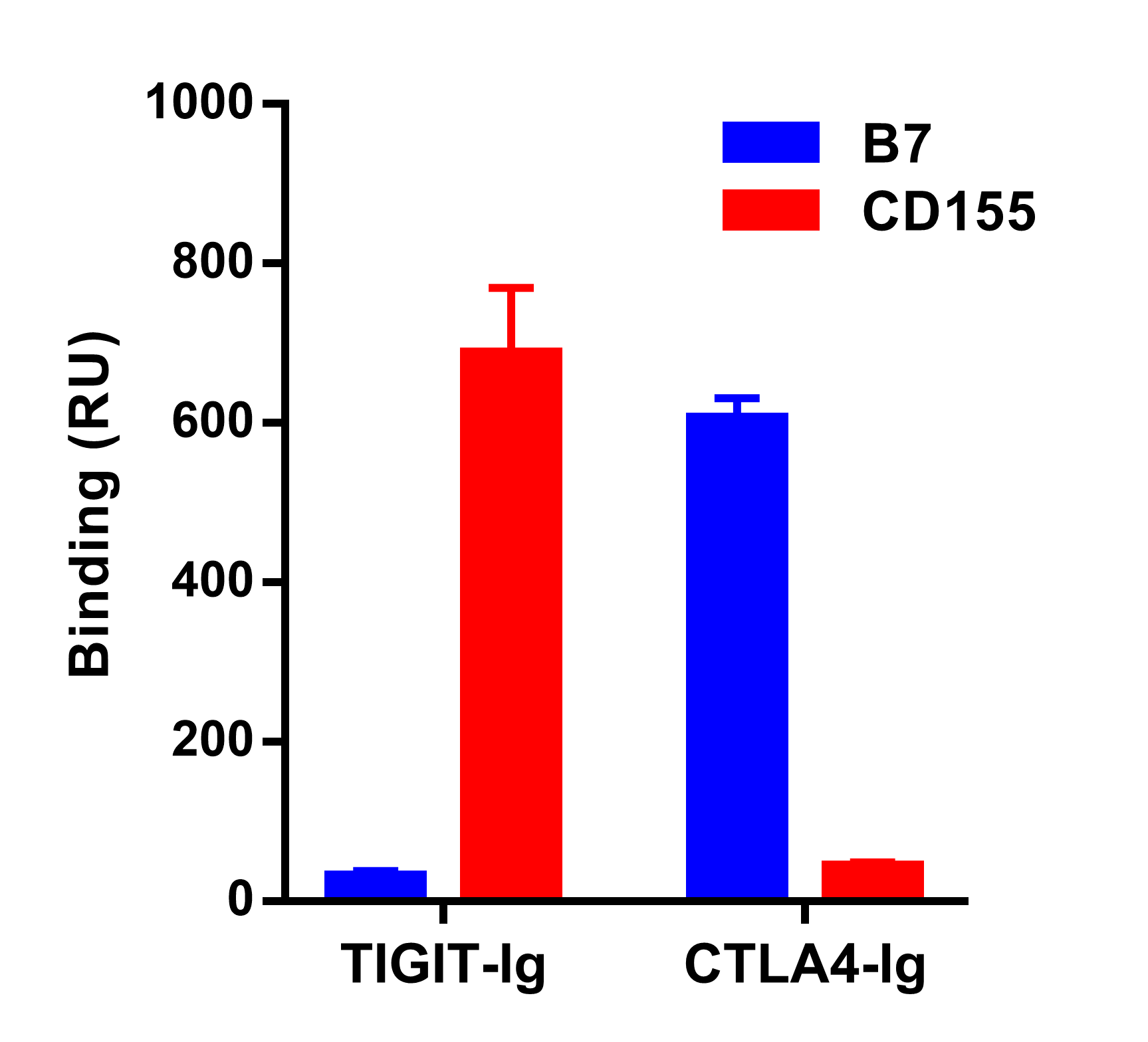

#### **Figure S1. Binding of Fusion proteins to CD155 and B7.**

Fusion proteins was tested for binding to immobilized CD155 or B7 protein using surface plasmon resonance on a BIAcore 2000 instrument. Binding was quantified as an increase in RU at 60 s after the end of injection compared with a baseline established 20 s before injection.

### Supplementary Tables

#### Table S1. Selected analytical data and pharmacokinetic parameters of recombinant fusion proteins in mice.

| Parameter^a^ | CTLA4-Ig | TIGHT-Ig |
| --- | --- | --- |
| HMW formation after storage(% SEC area)^b^ | < 0.1 | < 0.1 |
| LMW formation after storage(% SEC area)^b^ | < 0.1 | < 0.1 |
| AUC (day μg ml^-1^) | 470.7 | 555.1 |
| *T_1/2_(*day) | 7.4 | 7.9 |
| CL (ml day^-1^ kg^-1^) | 7.6 | 5.9 |
| VSS(ml kg^-1^) | 89.0 | 78.1 |

^a^ Pharmacokinetic parameters were calculated using a noncompartmental analysis. AUC, area under the concentration versus time curve; t1/2, half-life; CL, clearance; VSS, steady-state volume of distribution.

^b^ Quiescent storage for 3 wk, 40 °C, 1 mg/mL

#### Table S2. Biochemical characteristics after 10 weeks of treatment

| Parameter^a^ | CTRL-Ig | TIGHT-Ig |
| --- | --- | --- |
| BW (g) | 24.5 ± 0.52 | 24.8 ± 0.72 |
| BUN (mg/dL) | 30.31 ± 1.31 | 25.44 ± 0.64* |
| Cre (mg/dL) | 0.34 ± 0.02 | 0.20 ± 0.03** |
| HDL/LDL | 7.46 ± 0.54 | 8.33 ± 0.39 |
| TG (mg/dL) | 79.51 ± 5.88 | 63.67 ± 3.55* |
| TC (mg/dL) | 125.94 ± 4.37 | 95.81 ± 6.54* |
| BP (mmHg) | 100.5 ± 0.94 | 94.9 ± 1.52* |
| UACR (mg/mg) | 99.8 ± 4.37 | 55.1 ± 3.11** |

Data are presented as mean ± SD; *p < 0.05, and **p < 0.01, significant difference compared to the control group determined by Student's t test. BP: blood pressure; BUN: blood urine nitrogen; BW: body weight; Cre: creatinine; HDL: high-density lipoprotein; LDL: low-density lipoprotein;

TC: total cholesterol; TG: triacylglycerol; UACR: urinary albumin/urinary creatinine.
